# Fenretinide targets GATA1 to induce cytotoxicity in GATA1 positive Acute Erythroid and Acute Megakaryoblastic Leukemic cells

**DOI:** 10.1101/2025.01.19.633759

**Authors:** Yasharah Raza, Manal Elmasry, GuiQin Yu, Sam Blake Chiappone, Suhu Liu, Chiara Luberto

## Abstract

Patients with Acute Myeloid Leukemia (AML) subtypes, acute erythroleukemia and acute megakaryocytic leukemia (M6 and M7 AMLs, respectively) have a median survival of only a few months with no targeted effective treatment. Our gene expression analysis using the Cancer Cell Line Encyclopedia and CRISPR screen from DepMap showed that M6/M7 AMLs have high levels of the transcription factor *GATA1* and depend on GATA1 for survival. While GATA1 was shown to support AML cell proliferation and resistance to chemotherapy, GATA1 has long been considered “undruggable”. Here, we identify the small molecule N-(4-hydroxyphenyl)retinamide (4-HPR, Fenretinide) as a novel GATA1 targeting agent in M6 and M7 AML cells, with nM to low μM concentrations of 4-HPR causing loss of GATA1. In M6 AML OCIM1 cells, knock-down of *GATA1* induced cytotoxicity similarly to low doses 4-HPR while overexpression of GATA1 significantly protected cells from 4-HPR-induced cytotoxicity. In M6 AML cells resistant to current standard-of-care (SOC) Azacytidine plus Venetoclax, 4-HPR synergized with SOC overcoming cell resistance to the drugs. As single-agent, 4-HPR outperformed SOC. In M6 AML cells sensitive to SOC, 4-HPR enhanced and prolonged the growth inhibitory effect of SOC. 4-HPR is a synthetic derivative of vitamin A, and numerous clinical trials have supported its safe profile in cancer patients; therefore, targeted use of 4-HPR against M6 and M7 AMLs may represent a novel therapeutic window.

**Key Points:**

- Fenretinide (4-HPR) targets the transcription factor GATA1, which was previously thought to be “undruggable” and induces GATA1 loss.
- M6 and M7 Acute Myeloid Leukemias (AML) have enriched expression of GATA1 and they can be considered GATA1 positive.
- Loss of GATA1 contributes significantly to 4-HPR cytotoxicity in M6 OCIM1 cells.
- 4-HPR treatment overcomes chemotherapeutic resistance in M6 Acute Myeloid Leukemia cells, synergizes with standard-of-care and outperforms standard-of-care as a single agent.

## I. INTRODUCTION

Acute myeloid leukemia (AML) is a cancer of the blood and bone marrow characterized by aberrantly differentiated and out-of-control proliferating cells. According to the French, American, British classification of AML subtypes, which grouped neoplasms based on specific morphological/functional characteristics of tumor cells, M6 and M7 AMLs (also known as acute erythroleukemia and acute megakaryocytic leukemia, respectively) are very rare subtypes of AML. A revised 2022 classification of blood neoplasms by the World Health Organization based on molecular characteristics still defines these neoplasms as distinct entities(1). AML has an incidence of 4.1 cases per 100,000 people in the United States annually(2). M6 and M7 AMLs together represent approximately 7% of all AMLs (represented by 5% acute erythroleukemias, less than 1% Pure Erythroid Leukemias and 1% megakaryocytic leukemias)(3–7). Among all AMLs, M6 and M7 AMLs are the most lethal, exhibiting a poor response to standard therapy due to chemotoxicity and resistance (8, 9). The average survival has been reported as low as 3 to 9 months after diagnosis, and studies cite a median survival of M6 AML patients as merely 1.8 months(10).

Even in the current era of multiple novel therapies introduced for other malignancies, there is no effective therapy to offer to M6 and M7 AML patients (7, 10–12), and these high-risk patients are treated with a combination of Azacytidine (hypomethylating agent; Aza) and Venetoclax (BH3 mimetic inhibitor of BCL2; Veneto) (13). While this combination is effective in other subtypes of AML, it has not improved the survival of M6/M7 AML patients(10), and in fact, M6/M7 AML leukemic cells exhibit resistance (14). Additionally, while Veneto-based treatments are considered low intensity, they have been found to induce deep neutropenia, which increases vulnerability to infection (15, 16). Therefore, in addition to failing to improve survival of M6/M7 AML patients, the current standard-of-care (SOC) puts already high-risk patients at further increased risk of infection, requiring them to use additional drugs to mitigate this risk. Patients taking SOC are also at an increased risk for other adverse hematologic side effects such as thrombocytopenia, leukopenia, and anemia (17). Thus, there is a desperate need for novel targets and therapeutic strategies for these AMLs.

Our analysis of TCGA and BEAT AML 2.0 data indicates that M6 and M7 AML patient cells present high expression of the transcription factor, *GATA1*. In normal physiological conditions, GATA1 functions as the master regulator of erythroid and megakaryocytic differentiation (18). However, evidence in AML erythroblasts also implicate GATA1 in supporting their glycolytic metabolism, proliferation, and resistance to chemotherapy (19–21). Transcription factors, like GATA1, have long been considered “undruggable”. While this is beginning to change with emerging therapies directed to modulate gene expression, targeting transcription factors still remains a challenging therapeutic strategy.

We have identified N-(4-hydroxyphenyl)retinamide (4-HPR or Fenretinide) as a novel GATA-targeting agent in M6/M7 AML cells that can overcome chemotherapeutic resistance, synergize with the current SOC, and out-perform the SOC as a single agent. 4-HPR is a synthetic derivative of retinoic acid with anti-cancer effects in cell culture and preclinical mouse models for a variety of cancers (22–25). In animal models and Phase I and II clinical trials, 4-HPR has already shown a favorable toxicity profile (26–30). Therefore, our breakthrough results support the consideration of 4-HPR as a promising new targeted therapeutic option for patients with M6/M7 AML, warranting further pre-clinical investigation.

## **II.** MATERIALS & METHODS

### Cell Lines

HL-60, TF-1, and Hel 92.1.7 cells were purchased from the American Type Culture Collection (ATCC). OCIM1, Hel, and MOLM-16 cells were obtained from Deutsche Sammlung von Mikroorganismen und Zellkulturen (DSMZ, Brunswick, Germany). Hel and Hel92.1.7 were grown in RPMI-1640 (Life Technologies Corporation, Carlsbad, CA) with 10% HI-FBS (Heat Inactivated-Fetal Bovine Serum, Life Technologies Corporation, Carlsbad, CA). TF-1 were grown in RPMI-1640 with 10% HI-FBS and 2ng/ml GM-CSF (Stem Cell Technologies; Cat No. 78015). MOLM-16 were grown in RPMI with 20% HI-FBS. OCIM1, and HL-60 cells were grown in IMDM with 20% HI-FBS. Growth media also contained 100U/mL Penicillin and 100μg/mL Streptomycin (Life Technologies Corporation, Carlsbad, CA). Cells were routinely checked for mycoplasma contamination with the Lonza mycoAlert detection kit (Morristown, NJ, USA).

### Drug Treatments

Fenretinide (4-HPR; Sigma-Aldrich, Cat No. 65646-68-6) stock solution was prepared in ethanol. Venetoclax (ABT-199; Cat No. A12500; Adooq Bioscience) and 5-Azacytidine (Aza; Cell Signaling Technology; Cat No. 50646) stock solution was prepared in DMSO. At the time of treatment, cells were seeded at a concentration of 0.2×10^6^ cells/mL and incubated for up to 48 hours. 4-HPR treatment following *GATA1* knockdown were initiated 14 hours post-electroporation.

### Measuring Drug Synergy

OCIM1 cells were given doses of Aza, Veneto, and 4-HPR according to clinically relevant blood plasma levels, as previously reported (31–34). Doses for 4-HPR were 0, 0.05, 0.1, 0.175, 0.25, 0.35, 0.5, 1, and 2μM and doses for Veneto were 1, 2, or 3μM. Aza treatments were based on kinetics of plasma concentrations (31); cells were treated with 0, 2, 5, or 7μM Aza for 30 minutes, washed and then treated for the remainder of time with 0 or 0.5μM Aza in addition to combinations of 4-HPR and/or Venetoclax as indicated. Viability was measured at 48h by MTT assay. Synergy scores were calculated according to the Bliss independence score (35). Synergy scores greater than 10 are considered synergistic.

### *GATA1* Knockdown

The Neon Electroporation system (Life Technologies Corporation, Carlsbad, CA) was utilized for transfections. For each transfection, 2×10^6^ OCIM1 cells in log-growth phase were centrifuged at room temperature at a speed of 200 x g, for 5 minutes, and washed with 1x PBS (Life Technologies Corporation, Carlsbad, CA). Cell pellets were resuspended in 100μL Buffer-R and 20nM siRNA diluted in Buffer R was added (GATA1 siRNA: sc-29330, Santa Cruz Biotechnologies, Santa Cruz, CA or All-Star SCR siRNA as a negative control, Qiagen, Germantown, MD). Cells were electroporated (1000V, 50ms, 1 pulse) and transferred to a flask with 2mL complete media (pre-warmed). After 1 hour, 8ml of complete media were added (final growth conditions of 0.2×10^6^ cells/mL). Concentration of siRNA was optimized in preliminary experiments and calculated considering the final 10ml volume of medium.

### Stable Overexpression

#### *GATA1* over-expression construct

Human *GATA1* was PCR-amplified from pSIN4-EF1a-GATA1-IRES-Puro plasmid (Addgene plasmid # 61062; http://n2t.net/addgene:61062; RRID:Addgene_61062) (36) using forward and reverse primers as indicated in **Supplementary Table 1**. Amplified *GATA1* DNA and the destination vector pEF6/V5-His-TOPO (Invitrogen) were digested using Kpn-I and Xba-I restriction enzymes. Following digestion and gel extraction, the *GATA1* coding DNA sequence was ligated with the destination vector and insertion was confirmed by sequencing (**Supplementary Table 2)**. pEF6/*GATA1* and empty vector plasmids were prepared for transfections using the Endofree Plasmid Maxi Kit (Qiagen, Valencia, CA) as per manufacturer’s instructions. 10μg DNA was used for electroporation as described above (1000V, 50ms, 1 pulse). Cells were maintained in selection with Blasticidin (Life Technologies Corporation; Ref # A11139-03; Grand Island, NY) at 30μg/mL.

### Western Blotting

Cells (2-3 million) in 125-200μL of 1% SDS (Sodium Dodecyl Sulphate) were tip-sonicated (3x for 15s each at power output 3; Sonifier Cell Disruptor Model W185 (Plainview, NY)). After addition of 6x loading buffer, cell lysates were boiled (7 minutes). Forty μg of proteins were loaded in an SDS-PAGE (polyacrylamide gel electrophoresis) gel, run, and transferred onto nitrocellulose membranes at 80V. Membranes were blotted with primary antibodies against either GATA1 (N1 rat monoclonal, 1:1000 dilution, sc-266, Santa Cruz, Dallas, TX) or β-actin (1-19 goat polyclonal, 1:2500 dilution, sc-1616, Santa Cruz Biotechnology, CA). Secondary antibodies (1:5000; Jackson ImmunoResearch Laboratories, West Grove, PA) were diluted in 5% milk in PBS-T accordingly. Band intensities in western blots were quantified using Adobe Photoshop 2023 and normalized against actin.

### RNA extraction

1-2×10^6^ cells were used to extract RNA with the RNeasy mini kit (Qiagen, Germantown, MD). Cells were centrifuged at 1000 x g at 4°C for 5 minutes. Cells were then resuspended in lysis buffer with fresh β-mercaptoethanol (Sigma-Aldrich, St. Louis, MO), and RNA and cDNA were prepared as previously described (37).

### Quantitative Real-Time PCR (qRT-PCR)

qRT-PCR was performed using the Applied Biosystem, 7500 Real time PCR system. Results were analyzed using mean of normalized expression (MNE) (38). Taqman Probes for *GATA1* (Hs01085823_m1) and *18S* (Hs99999901_s1) (Thermo Fisher Scientific, Waltham, MA) were used with the Biorad iTaq Universal Supermix (Bio-Rad, Hercules, CA, USA), according to manufacturer’s instructions.

### Cell Viability Assays

#### Trypan Blue Exclusion assay

A 1:1.5 dilution of trypan blue (Life Technologies, CA) and cells was prepared, and immediately counted manually in a hemocytometer. Blue stained cells were considered dead.

#### Lactate dehydrogenase (LDH) assay

LDH activity was measured utilizing an LDH Assay Kit (Abcam; ab65393) according to the manufacturer’s protocol with minor modifications. For total LDH: 200ul of 2.5% Triton-X were added to 0.5-1mL cell culture, vortexed, incubated at room temperature for 30 minutes, and spun down at 800 x g to collect supernatant. For released LDH, 200μL of fresh medium was added to the 0.5-1mL cell culture (to account for the Triton added for the total LDH), cell suspension was then centrifuged at 800 x g for 5 minutes, and medium was collected. For all samples (lysates and media), 2μL were added to 100μL reaction mix (100:2; LDH Assay Buffer:WST substrate mix) in a 96-well plate. After 30 minutes incubation at 37°C, absorbance was measured at 450nm with a reference wavelength of 650nm.

#### MTT (3-(4,5-dimethylthiazol-2-yl)-2,5-diphenyltetrazolium bromide) tetrazolium reduction assay

The MTT assay was performed as previously described for suspension cells, with some modifications (39). 100μL of cell suspension was added to 25μL MTT substrate and incubated for 2 hours in a 96-well plate. MTT substrate was prepared in 1xPBS at 5mg/mL, sterile-filtered and stored at 4°C, protected from light. Then, 125μL MTT solubilization solution (10% SDS aqueous solution in 0.01 N HCl) was added to each well and incubated overnight at 37°C. Absorbance was measured at 590nm.

### Statistics

All analyses of significance were conducted with Prism 9 (GraphPad, La Jolla, CA) based on at least 3 biological replicates, unless otherwise indicated. Significance was determined using unpaired Student’s t-test or Two-ways ANOVA with post hoc Tukey’s multiple comparisons test.

### Analysis of Publicly Available Datasets

Gene expression profiles of AML cell lines were visualized using DepMap Portal (Broad Institute), utilizing data from the Cancer Cell Line Encyclopedia. *GATA1* dependency was assessed using DepMap Portal (Broad Institute) with data from *GATA1* Gene Effect (Chronos) CRISPR (DepMap Public 22Q4+Score Chronos) represented for AML cell lines.

For analysis of GATA1 Target Pathways, data from the ENCODE project was downloaded from the Gene Expression Omnibus (GEO) for fetal Peripheral Blood Derived Erythroblast (PBDE) and K562 cells (GSM935333 and GSM935540, respectively) as narrowPeak files. These files were analyzed by EaSeq(40). Briefly, GATA1 peaks located within 1 kilobase upstream of the start and downstream of the end of a gene were considered binding sites for GATA1 on that gene, as these could be binding sites within the promoter (transcriptional start site-proximal peaks) or enhancers (other peaks). The genes that had a GATA1 peak were then analyzed for gene ontology using the web-based program EnrichR and the Kyoto Encyclopedia of Genes and Genomes (KEGG) database of biological pathways. The negative of the logarithm base 10 of each pathway’s p-value was then calculated and plotted.

## **III.** RESULTS

### M6 and M7 AML have the worst prognosis of all AML subtypes and rely on high expression of *GATA1*

Old age is an unfavorable prognostic factor for AML patients (41), and while patients with M6 and M7 AML tend to be diagnosed at old age (median ages 60.3 years for M6 AML patients and 71 years for M7 AML patients) (**Figure 1A**), additional factors also contribute to their dire prognosis. In fact, analysis of TCGA survival data for age-matched AML patients (age equal or greater than 60 years at time of initial diagnosis) shows zero percent probability of survival for M6 and M7 AML patients past 215 days (about 7 months) and 427 days (about 14 months) after diagnosis, respectively (**Figure 1B**). This outcome is worse than any other AML subtype. Therefore, M6 and M7 AML are inherently more aggressive than the other AML subtypes. One characteristic of these M6 and M7 AML is their resistance to current therapies, and one potential modulator of this chemoresistance is the transcription factor GATA1. While GATA1 has been shown to contribute to leukemic cell survival and chemotherapeutic resistance (19–21), the level of expression of *GATA1* has not been thoroughly investigated in M6 and M7 AMLs. Therefore, we evaluated *GATA1* expression in these AMLs using the Cancer Cell Line Encyclopedia (CCLE - an open access repository for genomic annotations, gene expression and pharmacological characterization for a large number of cancer cell lines) which confirmed *GATA1* positivity in all acute erythroleukemic (M6) and megakaryocytic (M7) cell lines (**Figure 1C**). Importantly, analysis of CRISPR/Cas9 mediated knockout of *GATA1,* as reported by DepMap (an open-access portal that provides information on gene dependency of various cell lines), showed that M6 and M7 AML cell lines are highly dependent on *GATA1* (more negative CRISPR gene effect) compared to other subtypes of AML **(Figure 1D).** This is also in agreement with results reported in a recent report (14).

**Figure 1:**
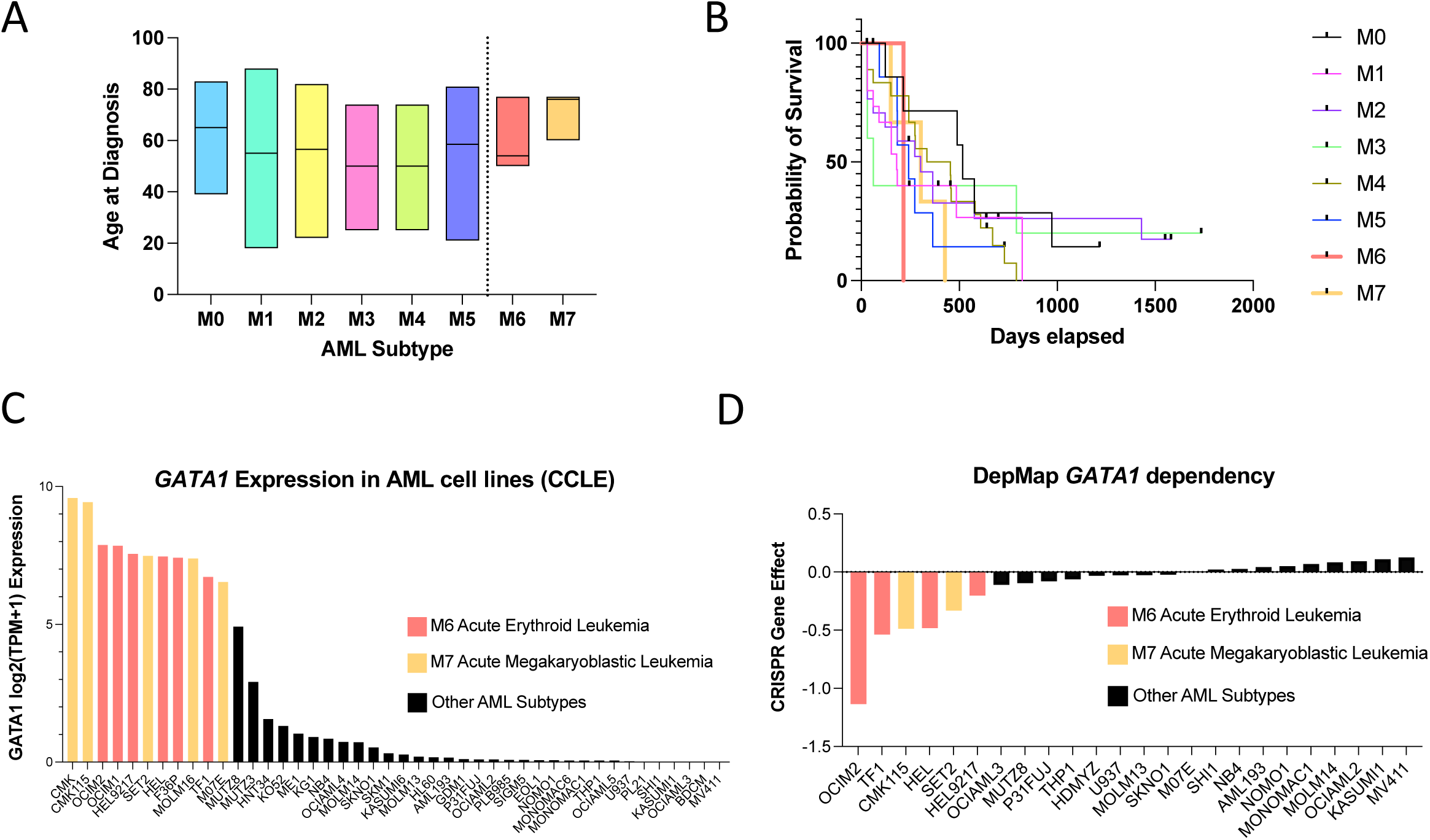
M6 and M7 AML have the worst prognosis of all AML subtypes and rely on high expression of *GATA1*. (**A**) Age at diagnosis across AML subtypes according to data from The Cancer Genome Atlas (TCGA) LAML dataset. (**B**) Probability of 5-years survival for patients 60 years and older, after AML diagnosis across AML subtypes according to the TCGA LAML dataset. (**C**) *GATA1* expression across AML cell lines reveal high GATA1 in M6 and M7 AML cells, according to data from Cancer Cell Line Encyclopedia (CCLE). (**D**) DepMap GATA1 dependency according to the GATA1 Gene Effect (Chronos) CRISPR (DepMap Public 22Q4+Score Chronos) represented for AML cell lines.

Altogether, these results suggest that *GATA1* expression is a signature of M6 and M7 AMLs and contributes to M6 and M7 AML cell survival.

### N-(4-hydroxyphenyl)retinamide (Fenretinide or 4-HPR) is a novel GATA1 targeting agent and induces cytotoxic and cytostatic effects in M6 and M7 AML cells

The induction of GATA1 in normal hematopoietic progenitors is essential for erythropoiesis (42–45). On the other hand, high GATA1 level in all M6/M7 AML cells (**Figure 1C**) and in a large fraction of Chronic Myelogenous Leukemia (CML) cell lines (**Figure S1**) is not sufficient to induce erythroid differentiation and it might even sustain proliferation and promote chemotherapeutic resistance as was shown in blasts from patients with acute megakaryocytic leukemia treated with Cytarabine and Daunorubicin (20). Interestingly, by probing publicly available ChIPSeq datasets, we compared GATA1 gene targets in fetal peripheral blood-derived erythroblasts (PBDEs) to those in K562, a GATA1-positive CML erythroleukemic cell line (**Figure S1**) and notable differences were found (**Figure 2**). Using EASeq and the Kyoto Encyclopedia of Genes and Genomes (KEGG) pathways, porphyrin/chlorophyll metabolism and autophagy were identified as the top two most significant pathways targeted by GATA1 in fetal PBDEs (**Figure 2A**); targets in these pathways are important for heme biosynthesis (46) and elimination/recycling of organelles as occurs in terminal erythroid differentiation (47), respectively. On the other hand, in K562 erythroleukemic cells, the top hits are pathways of sphingolipid signaling and sphingolipid metabolism (**Figure 2B**). This suggests that the gene expression program instigated by GATA1 may be differently wired in erythroleukemia versus normal erythroblasts with sphingolipid signaling being a prime GATA1 target pathway in the first. Analysis with EnrichR confirmed the identified GATA1-target genes (**Figures 2A** and **B**) as *bona fide* GATA1 targets (**Figures 2C** and **D**).

**Figure 2:**
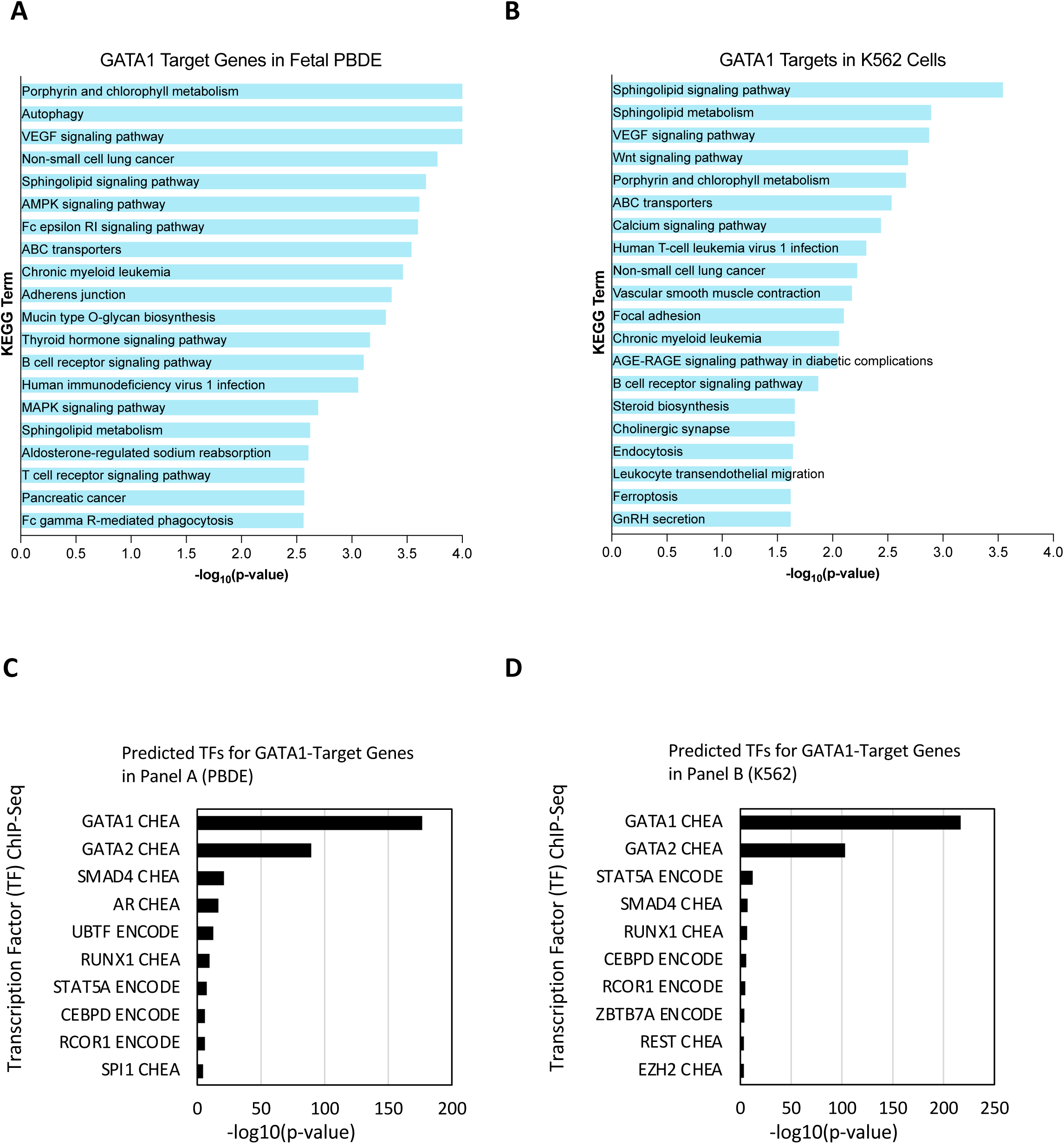
The representation of the pathways regulated by GATA1 gene targets is different between normal fetal peripheral blood-derived erythroblasts (PBDE) and erythroleukemic K562 cells. Kyoto Encyclopedia of Genes and Genomes (KEGG) pathway analysis of GATA1 target genes in (**A**) fetal peripheral blood-derived erythroblasts (PBDE), and (**B**) erythroleukemic K562 cells (GATA1^+^ erythroleukemic cells). Significance is quantified as the -log_10_(p-value) of gene ontology enrichment terms exhibiting genes with a detected GATA1 peak within 1000bp. To confirm accuracy of analysis, transcription factors (TFs) for GATA1-gene targets in fetal PBDEs (**C**) and K562 cells (**D**) are predicted by EnrichR and graphed with -log_10_(p-value) quantifying significance.

Among the GATA1 sphingolipid gene targets in K562 cells is the gene encoding for the dihydroceramide desaturase 1 (*DEGS1*). Notably, the small molecule N-(4-hydroxyphenyl)retinamide (Fenretinide or 4-HPR) is a DEGS1 inhibitor with reported pre-clinical anticancer activity and a safe profile in clinical trials (48–50); therefore we tested 4-HPR’s antileukemic effects in a panel of M6 and M7 AML cell lines (**Figure 3A-C**). As shown in the Figure, 4-HPR caused a significant decrease in live cell number (**Figure 3A**) and a dose-dependent increase in cytotoxicity measured by trypan blue positivity (**Figure 3B**) and percent of lactate dehydrogenase (LDH) released into the cell culture media (**Figure 3C**). The antileukemic effects of 4-HPR in these M6 and M7 AML cells after 48 hours of treatment were observed at concentrations equal or below 500nM in OCIM1 and TF-1 cells and equal or below 1.5μM for the other cell lines; on the other hand, 4-HPR treatment did not impact GATA1 negative HL-60 cells at doses up to 2μM (**Figure S2A**). While 4-HPR is a synthetic retinoid, its cytotoxic effect in OCIM1 M6 AML cells seems to be independent from activation of the retinoic acid receptor, as treatment with *all-trans* Retinoic Acid at concentrations up to 10μM did not affect cell number (**Figure S2B**).

**Figure 3:**
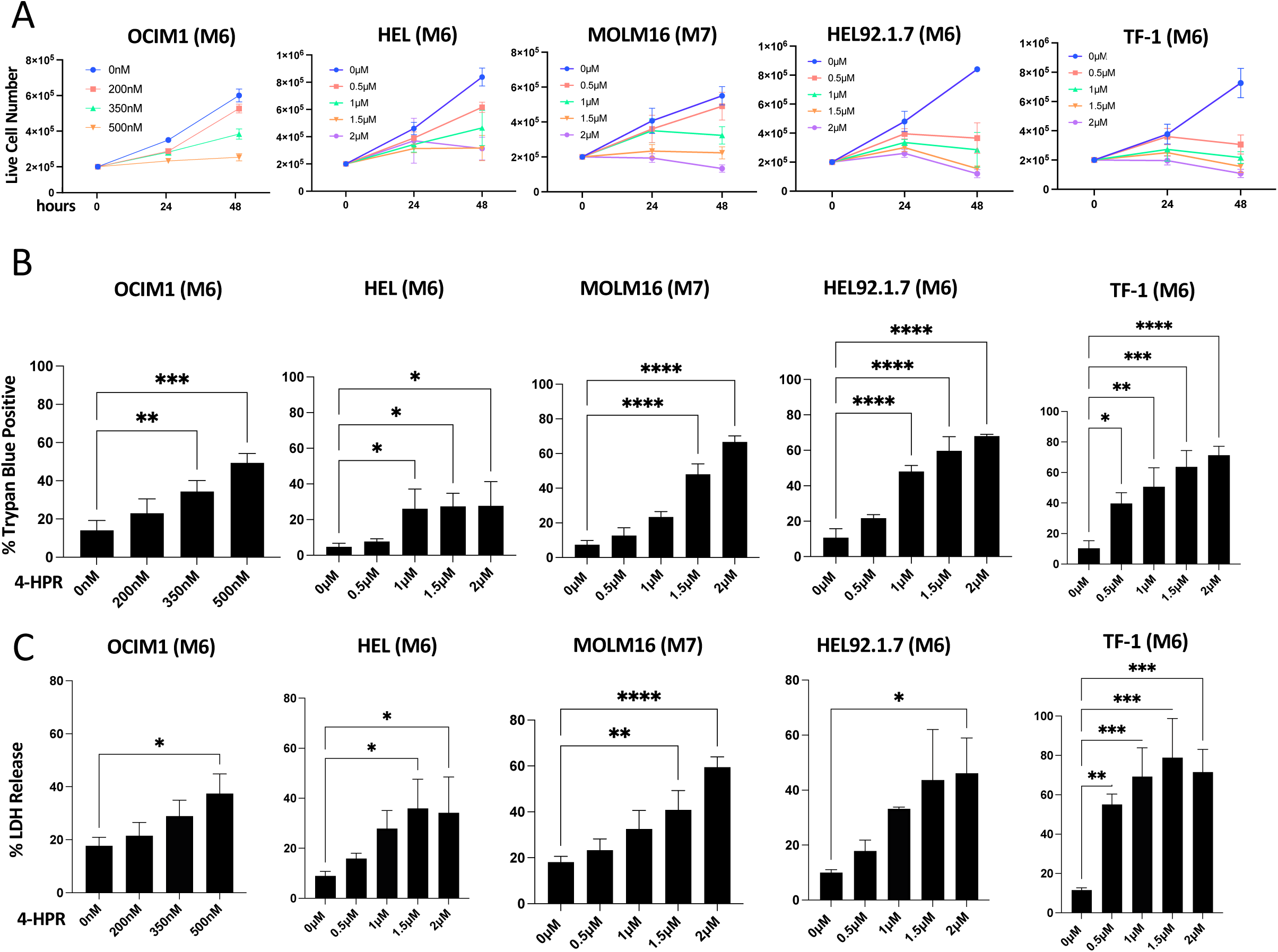
4-HPR Treatment of GATA1^+^ AML Cell Lines is cytotoxic/cytostatic. (**A**) 4-HPR treatment on live cell number in OCIM1, Hel, MOLM-16, HEL 92.1.7 and TF-1 cell lines. 4-HPR has a dose-dependent cytotoxic effect measured by trypan blue positivity (percentage of total cell number) (**B**) and by lactate dehydrogenase release (LDH; percentage of LDH released in the medium over total LDH) (**C**). Data are representative of three biological replicates; *p<0.05; **p<0.02; ***p<0.002, ****p<0.001

### GATA1 loss mediates the effects of Fenretinide in acute erythroleukemic M6 cell line OCIM1

Given the dependence of M6 and M7 AML cells on GATA1 (**Figure 1D**), we wondered whether 4-HPR affects GATA1. Intriguingly, 4-HPR targeted GATA1 protein causing its loss (**Figure 4A and B**). GATA1 loss was observed at the lowest concentrations of 4-HPR that caused significant cytostatic/cytotoxic effects (350nM in OCIM1 cells, 1μM in Hel cells, 1.5μM in MOLM-16 cells, 0.5μM in Hel 92.1.7 cells, and 0.5μM in TF-1 cells after 48 hours of treatment). These results support a novel targeting action of 4-HPR on GATA1 which parallels 4-HPR’s cytostatic/cytotoxic effects in M6/M7 AML cells.

**Figure 4:**
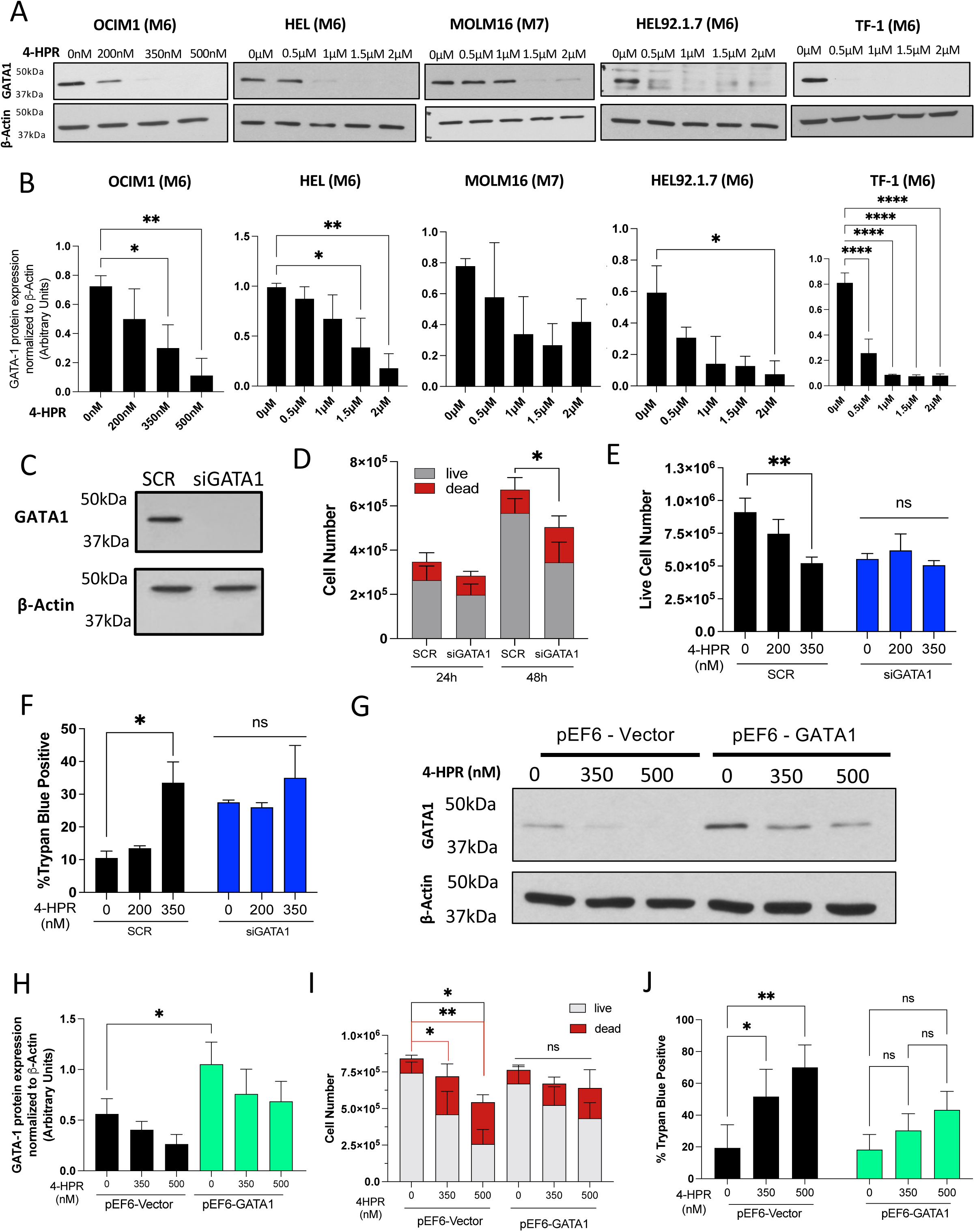
4-HPR targets GATA1 to mediate cytotoxic/cytostatic effects. (**A**) GATA1 protein expression levels by western blotting in different GATA1^+^ AML cell lines upon 4-HPR treatment. (**B**) Quantification of GATA1 levels upon 4-HPR treatment in respective GATA1^+^ AML cell lines. 4-HPR treatment leads to reduction of GATA1 levels. (**C**) siRNA targeting *GATA1* (siGATA1) leads to GATA1 protein loss as measured by western blotting; (**D**) siGATA1 causes cell death and decreases the cell number, with a significant decrease in live cell number between SCR and siGATA1; (**E**) the effect of siGATA1 is not additive to 4-HPR’s effect on live cell number, (**F**) or on trypan blue (TB) positivity (%TB positive). (**G**) GATA1 protein levels in OCIM1 cells stably overexpressing *GATA1* (pEF6-*GATA1*) and in vector control cells (pEF6-Vector) in absence (0nM) and presence of different concentrations of 4-HPR; (**H**) GATA1 protein quantification of E; (**I**) overexpression of GATA1 protects cells from 4-HPR’s effect on cell number. Red lines represent significance between dead cell number; (**J**) GATA1 overexpression protects cells from cytotoxicity (% trypan blue positivity). Data are representative of three biological replicates; *p<0.05; **p<0.02

To explore the contribution of GATA1 protein loss on cell proliferation and cell death after 4-HPR treatment, siRNA was used to downregulate GATA1 protein levels in OCIM1 M6 AML cells (**Figure 4C**). Downregulation of *GATA1* alone was associated with a significant decrease in live cell number (**Figure 4D**). Furthermore, treatment of *GATA1*-knockdown cells with 4-HPR at concentrations causing loss of GATA1 did not have a further impact on cell number or percent trypan blue positivity (**Figure 4E,F**). In OCIM1 cells, 4-HPR treatment reduced *GATA1* mRNA expression, suggesting the action of 4-HPR on pathways regulating *GATA1* expression (**Figure S3**).

As a complementary approach, we asked whether preventing GATA1 loss by its stable overexpression could protect from 4-HPR’s cytotoxic and cytostatic effects (**Figure 4G,H)**. GATA1 overexpression offered significant protection against 4-HPR-induced effects on cell number and cytotoxicity, measured by trypan blue positivity (**Figure 4I,J)**. Interestingly, 4-HPR was still partly effective in reducing levels of overexpressed GATA1 (**Figure 4G,H**) suggesting that, in addition to *GATA1* mRNA, 4-HPR also targets mechanisms regulating GATA1 protein. Altogether, these results suggest that GATA1 loss contributes to 4-HPR’s cytotoxic and cytostatic effects in OCIM1 cells.

### 4-HPR overcomes therapeutic resistance, synergizes with the current standard-of-care, and outperforms standard-of-care as a single agent

The standard therapeutic regimen for M6 and M7 AML patients consists of treatment with a hypomethylating (HMA) agent such as Azacytidine (Aza), in combination with Venetoclax (Veneto), a BCL-2 inhibitor (17). Notably, acute erythroleukemic cells were shown to be Veneto resistant (14), and patients have responded poorly and exhibited resistance against this treatment strategy (10). The cell line OCIM1 demonstrated high sensitivity to 4-HPR treatment, with effective doses in the nanomolar range (**Figure 3**) while also exhibiting resistance to Veneto (data not shown). Hence OCIM1 was used to assess the effect of 4-HPR in combination with the current standard-of-care (SOC). Cells were treated with a range of clinically relevant 4-HPR doses (0.05-2μM) which are below those found in plasma of patients who received 4-HPR orally or intravenously as part of past clinical studies (32, 33) (NSC374551), and the response was compared to combination of Aza plus Veneto. Additionally, 4-HPR doses were assessed for their ability to synergize with SOC. In patients, Aza is mostly administered intravenously (i.v.) at 75mg/m^2^ over 10-30 minutes once a day for 7 days. Aza’s plasma level increases very rapidly upon infusion and remains up for the duration of the treatment (for a 30 minutes infusion it reaches ∼5μM); once the infusion is terminated, the level of the drug drops immediately to ∼0.5-1μM and continues to quickly decrease for the next several hours (Australian Public Assessment Report for azacytidine) (31). Veneto is administered orally (at 400mg/day) and plasma concentrations are ∼3μM (34, 51). Thus, in accordance with clinical practice and reported drug plasma concentrations, we treated cells for 30 minutes with 0, 2, 5, or 7μM Aza, and then either incubated them with 0.5μM of Aza (with Aza maintenance) or with no Aza (without Aza maintenance) (**Figure 5**). Aza treatment was combined with either 0, 1, 2, or 3μM Veneto, and varied doses of 0.05-2μM 4-HPR for 48 hours and viability was measured via MTT assay.

**Figure 5:**
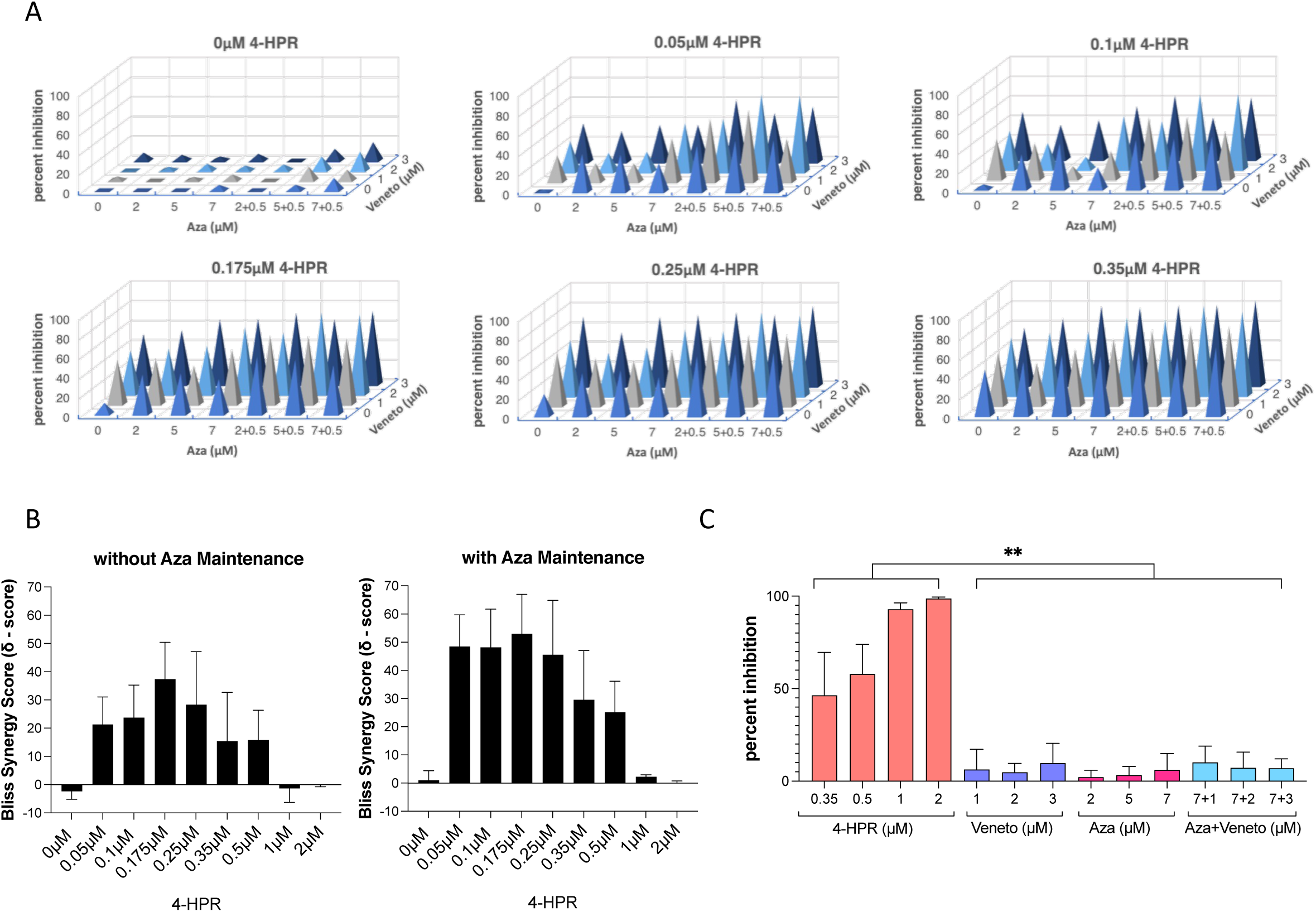
4-HPR treatment overcomes chemotherapeutic resistance, synergizes with the current SOC, and outperforms SOC as a monotherapy. OCIM1 cells were treated with or without 4-HPR, Aza and Veneto for 48 hours; viability was quantified by MTT assay. Reported is the percent inhibition of MTT absorbance considering cells treated with vehicle only as 100% viable. (**A**) The ability of 4-HPR to sensitize OCIM1 cells to cell death is represented by percent inhibition measured by MTT assay. Concentrations of Azacytidine (Aza) pre-treatment at 0, 2, 5, 7µM (with and without additional 0.5µM of Aza maintained in media) and Venetoclax (Veneto) at 0, 1, 2, and 3µM as indicated, were tested in the presence of 0, 0.05, 0.1, 0.175, 0.25, and 0.35µM of 4-HPR for sensitization to Aza+Veneto. (**B**) Synergy scores (δ – score) of 4-HPR (0, 0.05, 0.1, 0.175, 0.25, and 0.35µM) with Aza and Veneto, with or without maintenance of 0.5µM Aza in the media for 48 hours. δ – scores were determined by SynergyFinder (synergyfinder.fimm.fi), according to Bliss independence, where δ – score > 10 indicates synergism, and δ – score > -10 indicates antagonism. (**C**) 4-HPR as a single-agent on percent inhibition compared to Veneto and Aza monotherapies, and to Aza+Veneto combination therapy. Data are representative of the means of three biological replicates. **p<0.01. Significance is indicative of a summary of p-values. Each concentration of 4-HPR had a significant effect compared to each dose of Aza and/or Veneto.

In the absence of 4-HPR, OCIM1 cells’ viability remained greater than 80 percent (less than 20 percent cell inhibition) in all combinations of Aza+Veneto. Thus, without 4-HPR, OCIM1 cells are resistant to Aza, Veneto or Aza+Veneto (**Figure 5A**). Addition of 4-HPR to the therapeutic regimen overcame chemoresistance at doses as low as 0.05μM (**Figure 5A**). Not only 4-HPR overcame resistance to Aza and Veneto, but 4-HPR also strongly synergized with Aza and Veneto starting at 0.05μM, and continued to exhibit synergy through doses up to 0.5μM, according to the Bliss independence method of synergy determination, with synergy scores (δ – score) of 21.32, 23.74, 37.38, 28.35, 15.45, 15.78 for 0.05, 0.1, 0.175, 0.25, 0.35, and 0.5μM 4-HPR respectively (**Figure 5B, Figure S4**). This synergy is magnified when 0.5μM Aza is maintained for 48 hours of treatment, with average δ – scores of 48.45, 48.13, 52.99, 45.57, 29.59, 25.13 for 0.05, 0.1, 0.175, 0.25, 0.35, and 0.5μM 4-HPR respectively (**Figure 5B, Figure S4**). Higher concentrations of 4-HPR did not show synergy given the already significant cytotoxicity as single agent (**Figure S4**). In fact, 4-HPR outperformed Aza+Veneto as a single agent at 0.35μM, 0.5μM, 1μM, and 2μM, with an average inhibition of 46.38, 57.99, 92.86, 98.67 percent, respectively (**Figure 5C**). These results indicate potential for 4-HPR as a targeted monotherapy in these GATA1 positive AML cells.

### 4-HPR increases sensitivity to current SOC in cells already basally sensitive to the treatment and prolongs SOC’s efficacy

While M6 and M7 AML cells have been reported to be generally resistant to SOC, HEL and TF-1 cells showed some sensitivity to SOC compared to OCIM1 (**Figure 6**). However, cotreatment with 4-HPR at concentrations that decrease GATA1 levels (0.75μM for HEL and 0.35μM for TF-1) showed additive effect with SOC in both cell lines and, in the case of HEL, it significantly enhanced and prolonged the growth inhibitory effects of SOC, even when those seemed to subside as single treatments at the later time point. These results suggest that combination of SOC and 4-HPR produces beneficial effects even when cells might respond to SOC.

**Figure 6.**
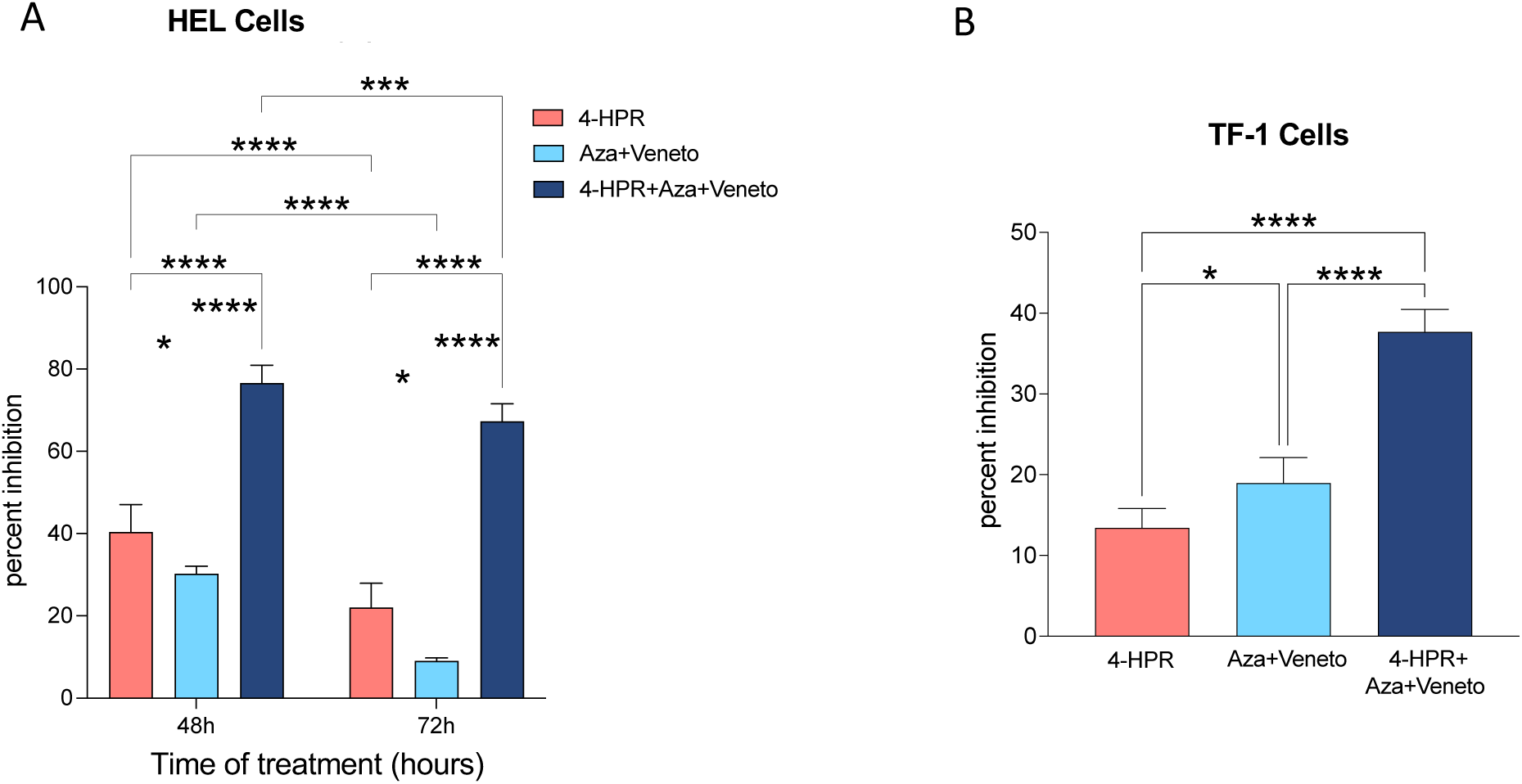
4-HPR increases sensitivity to current SOC in cells already basally sensitive to the treatment (HEL and TF-1) and prolongs SOC’s efficacy. HEL **(A)** and TF-1 **(B)** cells (0.2×10^6^ cells/ml) were treated with or without 4-HPR (0.75µM for HEL and 0.35µM for TF-1), Aza (maintenance protocol) and Veneto (3µM) for up to 48h (TF-1) or 72h (HEL). In case of Aza treatment, cells were exposed to 5µM Aza for 30min or Vehicle (DMSO) and, after a wash, cells were incubated with 0.1µM Aza or DMSO for maintenance. Viability was quantified by MTT assay and shown is the percent inhibition of MTT absorbance compared to vehicle control (considered 100% viable). Data are representative of the means of four biological replicates. *p<0.05; **p<0.01; ***p=0.0003; ****p<0.0001. Significance is indicative of a summary of p-values.

## **IV.** DISCUSSION

The present work demonstrates dependence of M6 and M7 AML cells on *GATA1*, it identifies 4-HPR as a novel modulator of GATA1, it shows that M6 and M7 AML cells are particularly sensitive to 4-HPR, and it demonstrates the ability of 4-HPR in sensitizing M6 and M7 AML cells to SOC and in outperforming it as a single agent.

Notably, 4-HPR has not been extensively investigated in clinical trials for blood cancers except for a Phase I trial in refractory and relapsed hematological malignancies (NCT00104923) (33). Given the low incidence of M6/M7 AMLs and the mixed typology of the patients included in the NCT00104923 trial, that study did not provide specific information regarding the potential therapeutic benefit of 4-HPR for M6/M7 AML. In other preclinical studies and in Phase I and II clinical trials, 4-HPR has shown a relatively safe profile with only some asymptomatic laboratory abnormalities (26–29). This is a critical consideration given GATA1’s role in normal erythropoiesis (52). However, despite the number of clinical trials performed with 4-HPR, therapeutic effectiveness of 4-HPR has been underwhelming, particularly for solid tumors, due, in large part, to the poor bioavailability of its formulations. New formulations of 4-HPR with better bioavailability have been reported (53), which combined with the low doses needed to induce GATA1 loss and anti-leukemic effects demonstrated in our studies, they represent a promising potential breakthrough in the treatment of M6 and M7 AML. Also relevant to the favorable effect of 4-HPR addition to sensitizing SOC, combination treatment with 4-HPR and Veneto has been already conducted in mice and tested on a model of neuroblastoma (54). High synergy and no toxicities were observed indicating that a combination of 4-HPR and Veneto is well tolerated.

Notably, all cell lines utilized in our studies have mutated TP53, indicating that 4-HPR does not require wild-type TP53 to target GATA1 and kill these cells. Historically, 4-HPR has been shown to induce apoptosis via p53-independent pathways and may have further efficacy in treating TP53 mutated AML beyond M6 and M7 AMLs (55). This is an important point since approximately 10-15% of all AML and approximately 30-35% of M6/M7 AMLs have mutated TP53 (56–59). Overall, AML patients with mutated TP53 have been shown to have poorer outcomes and exhibit chemotherapeutic resistance (56, 60–63). 4-HPR utility in AML patients with mutated TP53 is an exciting and promising avenue for further investigation, and at the very least, it would not constitute an obstacle to 4-HPR efficacy.

Recently, a critical and specific pro-survival function of BCL-XL was demonstrated for M6 and M7 AML cells as compared to other AML subtypes (14). Notably the expression of *BCL2L1,* the gene encoding for BCL-XL, is positively controlled by GATA1 (64, 65) and therefore it is possible that one of the mechanisms by which reduction of GATA1 levels affects survival of M6/M7 AML cells may be *via* altered expression of BCL-XL.

Mechanistically, we show that 4-HPR reduces *GATA1* mRNA by half; preliminary results using *GATA1* siRNA indicated that 50% down-regulation of its mRNA is enough to almost completely suppress GATA1 protein, implicating regulation of *GATA1* mRNA as a plausible mechanism of action of 4-HPR. This may not be the sole mechanism since overexpressed GATA1 originated from plasmid cDNA is also targeted by 4-HPR suggesting that multiple mechanisms (both at the gene and at the protein level) may be at work.

Additionally, we show that ATRA does not recapitulate the effects of 4-HPR on cell number, implying that 4-HPR is working on pathways independent from the retinoic acid receptor. This agrees with previous published literature (66–69). 4-HPR is known to have different targets through which it exerts its effects, and these targets/effects may be context dependent (24). For example, in ALL cells, 4-HPR has been found to target PKCδ, leading to increased levels of reactive oxygen species, which in turn induce autophagy and apoptosis (70). In hepatocellular carcinoma, 4-HPR treatment increases the expression of RARβ and initiates the nuclear export of the RARβ-Nur77 complex, which binds to Bcl-2 in the mitochondria causing Bcl-2 conformational change to promote apoptosis (71, 72). Interestingly, in our studies we find that loss of GATA1 by 4-HPR is independent from caspase activation as pretreatment with the pan caspase inhibitor Z-VAD did not prevent the decrease in GATA1 protein (data not shown). While this is in line with some published literature showing 4-HPR-induced cell death is both caspase dependent and independent (73), it is different from the caspase-3-dependent mechanism that drives the decrease of GATA1 necessary to complete the last phases of terminal erythropoiesis by committed erythroid precursors (74).

Importantly, 4-HPR also modulates sphingolipid metabolism via DEGS1 inhibition (48). We confirmed accumulation of dihydroceramide in HEL cells treated with 4-HPR (1.5μM, data not shown), however we were unable to fully downregulate *DEGS1* in tested M6/M7 cell lines and establish its potential active role in mediating 4-HPR-induced GATA1 loss. This will be fully investigated in future experiments.

### Data Sharing Statement

Requests for further information on reagents, methodologies and data should be directed and will be met by.

## Supporting information

Supplementary material all

## Acknowledgements

Yasharah Raza is a 2023-2024 American Fellow of the American Association for University Women and a recipient of a Turner Dissertation Fellowship by the Stony Brook University’s Center for Inclusive Education. This work was also supported by U.S. National Institutes of Health (NIH), National Cancer Institute Grant P01 CA097132 (to CL for Project #4), and the American Cancer Society – Institutional Research Grant # 21-143-01 and Stony Brook Cancer Center (to SL).

## Authorship

Y.R proposed the hypotheses, designed and performed the experiments, analyzed the data, prepared figures and wrote the manuscript; M.E. designed and performed the experiments, analyzed the data, prepared figures; S.B.C. analyzed the expression data, prepared figures and edited the manuscript; G.Y. performed experiments; S.L. proposed hypotheses, designed the study, analyzed the data, edited the manuscript; C.L. proposed the hypotheses, designed the study, analyzed the data and wrote the manuscript.

## Correspondence

Chiara Luberto, Department of Physiology and Biophysics, Stony Brook University; Stony Brook Cancer Center, MART Building, 9M-0828, 100 Nicolls Road, Stony Brook 11794, NY.

Suhu Liu, Division of Hematology & Oncology, Renaissance School of Medicine, Stony Brook University; Stony Brook Cancer Center; 100 Nicolls Road, Stony Brook 11794, NY;

---

**Figure S1:**
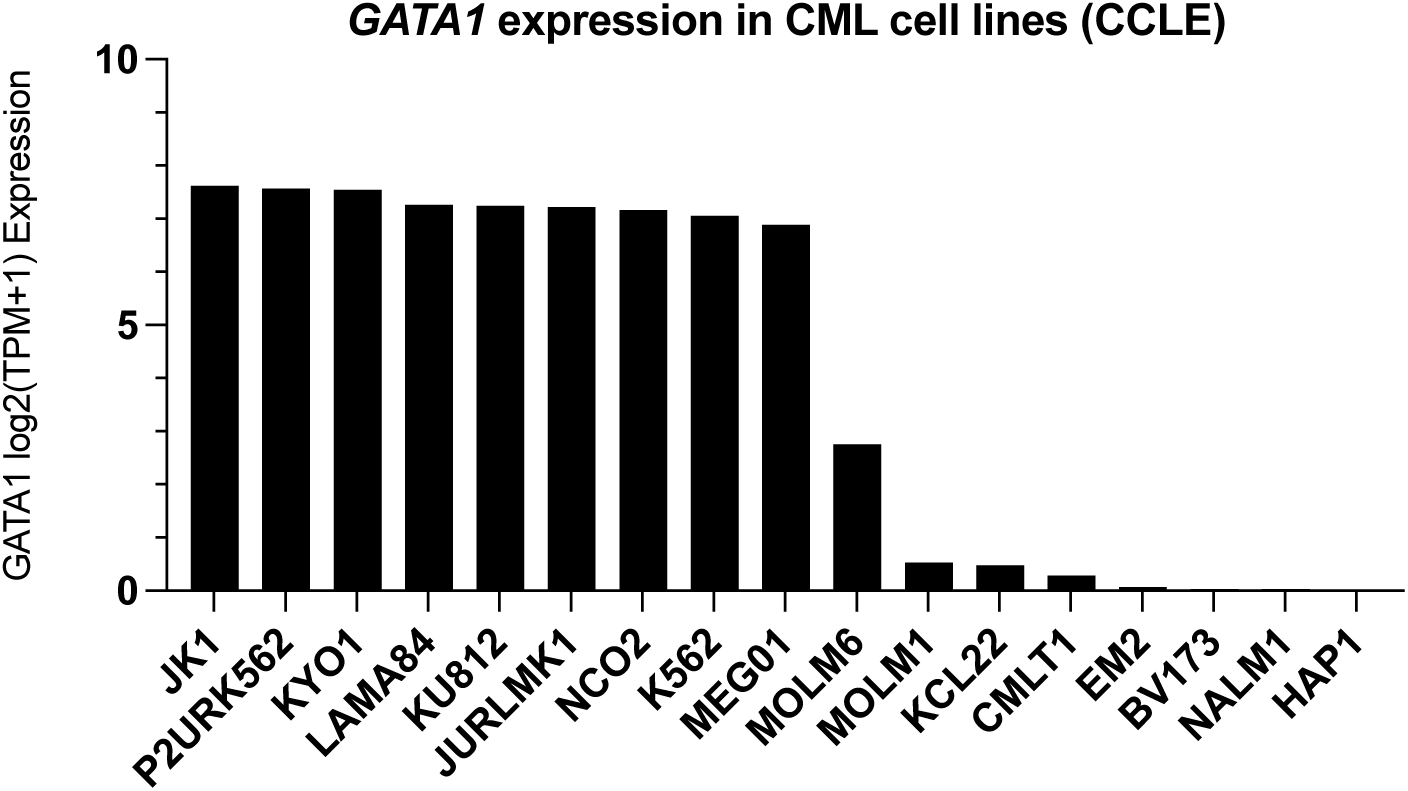
GATA1 expression in Chronic Myeloid Leukemia cell lines.

**Figure S2:**
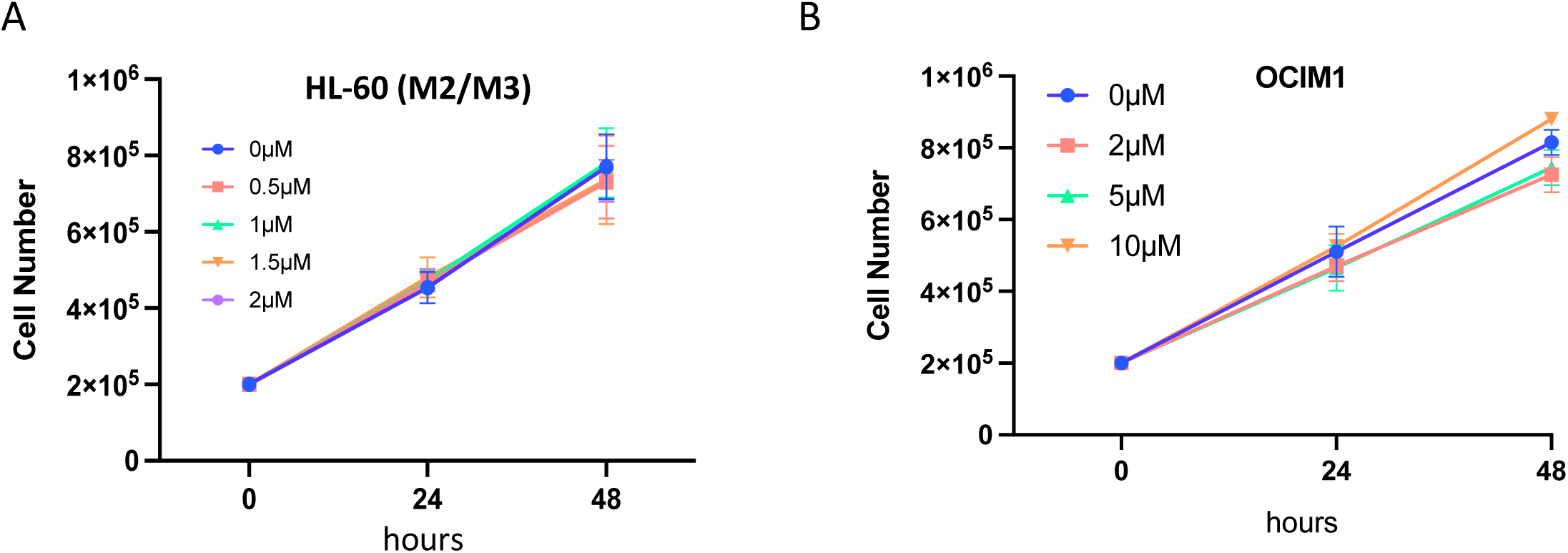
4-HPR treatment of GATA1 negative HL-60 cells and ATRA treatment of OCIM1 cells have no impact on cell number.

**Figure S3:**
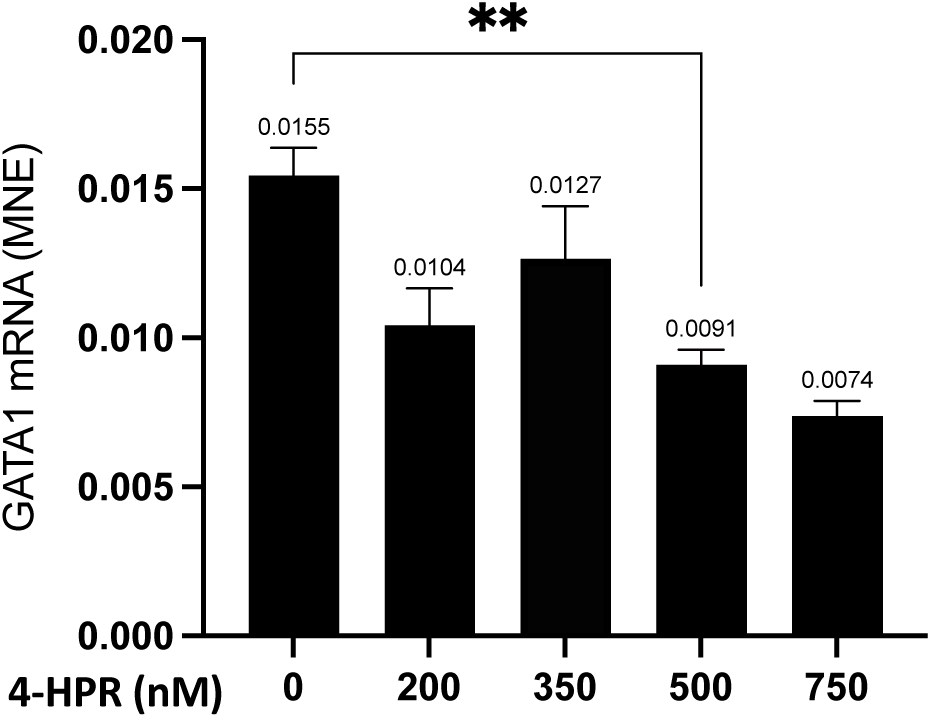
4-HPR treatment of OCIM1 cells reduces *GATA1* mRNA levels

**Figure S4:**
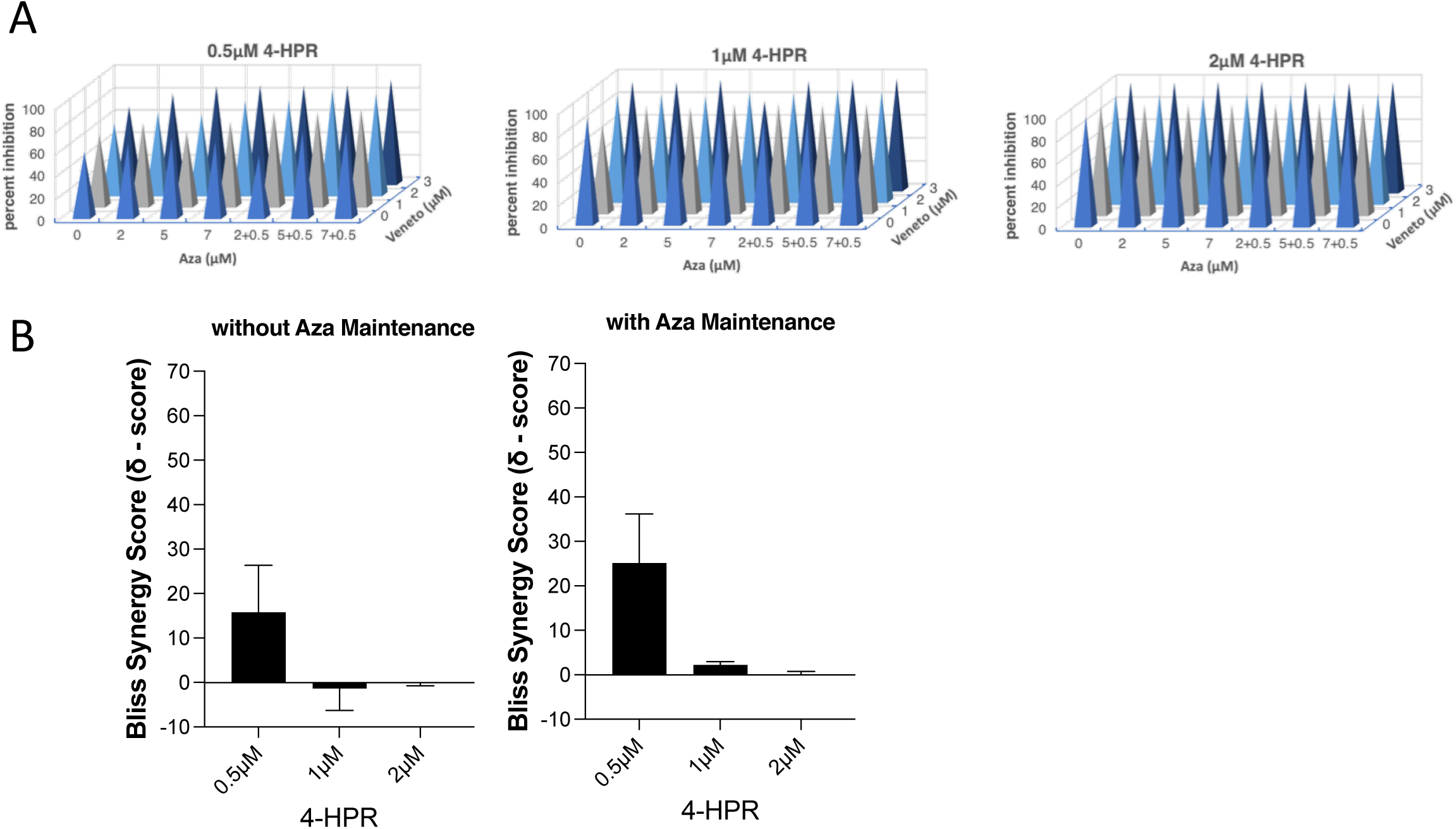
High doses of 4-HPR treatment in combination with Azacytidine and Venetoclax

## References

1. Khoury JD, Solary E, Abla O, Akkari Y, Alaggio R, Apperley JF, et al. The 5th edition of the World Health Organization Classification of Haematolymphoid Tumours: Myeloid and Histiocytic/Dendritic Neoplasms. Leukemia. 2022;36(7):1703–19.

2. Surveillance Research Program, Cancer Stat Facts: Leukemia — Acute Myeloid Leukemia (AML): National Cancer Institute SEER*Stat software (seer.cancer.gov/seerstat); [

3. Reinig EF, Greipp PT, Chiu A, Howard MT, Reichard KK. De novo pure erythroid leukemia: refining the clinicopathologic and cytogenetic characteristics of a rare entity. Mod Pathol. 2018;31(5):705–17.

4. Margolskee E, Mikita G, Rea B, Bagg A, Zuo Z, Sun Y, et al. A reevaluation of erythroid predominance in Acute Myeloid Leukemia using the updated WHO 2016 Criteria. Mod Pathol. 2018;31(6):873–80.

5. Duchayne E, Fenneteau O, Pages MP, Sainty D, Arnoulet C, Dastugue N, et al. Acute megakaryoblastic leukaemia: a national clinical and biological study of 53 adult and childhood cases by the Groupe Francais d’Hematologie Cellulaire (GFHC). Leuk Lymphoma. 2003;44(1):49–58.

6. Tallman MS, Neuberg D, Bennett JM, Francois CJ, Paietta E, Wiernik PH, et al. Acute megakaryocytic leukemia: the Eastern Cooperative Oncology Group experience. Blood. 2000;96(7):2405–11.

7. Wickrema A, Crispino JD. Erythroid and megakaryocytic transformation. Oncogene. 2007;26(47):6803–15.

8. Giri S, Pathak R, Prouet P, Li B, Martin MG. Acute megakaryocytic leukemia is associated with worse outcomes than other types of acute myeloid leukemia. Blood. 2014;124(25):3833–4.

9. Hasserjian RP, Zuo Z, Garcia C, Tang G, Kasyan A, Luthra R, et al. Acute erythroid leukemia: a reassessment using criteria refined in the 2008 WHO classification. Blood. 2010;115(10):1985–92.

10. Reichard KK, Tefferi A, Abdelmagid M, Orazi A, Alexandres C, Haack J, et al. Pure (acute) erythroid leukemia: morphology, immunophenotype, cytogenetics, mutations, treatment details, and survival data among 41 Mayo Clinic cases. Blood Cancer J. 2022;12(11):147.

11. Gera K, Martir D, Xue W, Wingard JR. Survival outcomes of acute erythroid leukemia in the United States: A 20-year population-based study. Journal of Clinical Oncology. 2023;41(16_suppl):e19031–e.

12. Tazi Y, Arango-Ossa JE, Zhou Y, Bernard E, Thomas I, Gilkes A, et al. Unified classification and risk-stratification in Acute Myeloid Leukemia. Nat Commun. 2022;13(1):4622.

13. Fleischmann M, Schnetzke U, Hochhaus A, Scholl S. Management of Acute Myeloid Leukemia: Current Treatment Options and Future Perspectives. Cancers (Basel). 2021;13(22).

14. Kuusanmaki H, Dufva O, Vaha-Koskela M, Leppa AM, Huuhtanen J, Vanttinen I, et al. Erythroid/megakaryocytic differentiation confers BCL-XL dependency and venetoclax resistance in acute myeloid leukemia. Blood. 2023;141(13):1610–25.

15. Samuels C, Abbott D, Niemiec S, Tobin J, Falco A, Halsema K, et al. Evaluation and associated risk factors for neutropenia with venetoclax and obinutuzumab in the treatment of chronic lymphocytic leukemia. Cancer Rep (Hoboken). 2022;5(5):e1505.

16. Waggoner M, Katsetos J, Thomas E, Galinsky I, Fox H. Practical Management of the Venetoclax-Treated Patient in Chronic Lymphocytic Leukemia and Acute Myeloid Leukemia. J Adv Pract Oncol. 2022;13(4):400–15.

17. DiNardo CD, Jonas BA, Pullarkat V, Thirman MJ, Garcia JS, Wei AH, et al. Azacitidine and Venetoclax in Previously Untreated Acute Myeloid Leukemia. N Engl J Med. 2020;383(7):617–29.

18. Ferreira R, Ohneda K, Yamamoto M, Philipsen S. GATA1 function, a paradigm for transcription factors in hematopoiesis. Mol Cell Biol. 2005;25(4):1215–27.

19. Kohrogi K, Hino S, Sakamoto A, Anan K, Takase R, Araki H, et al. LSD1 defines erythroleukemia metabolism by controlling the lineage-specific transcription factors GATA1 and C/EBPalpha. Blood Adv. 2021;5(9):2305–18.

20. Caldwell JT, Edwards H, Dombkowski AA, Buck SA, Matherly LH, Ge Y, et al. Overexpression of GATA1 confers resistance to chemotherapy in acute megakaryocytic Leukemia. PLoS One. 2013;8(7):e68601.

21. Bai Y, Qiu GR, Zhou F, Gong LY, Gao F, Sun KL. Overexpression of DICER1 induced by the upregulation of GATA1 contributes to the proliferation and apoptosis of leukemia cells. Int J Oncol. 2013;42(4):1317–24.

22. Zhang H, Xu H, Zhang R, Zhao X, Liang M, Wei F. Chemosensitization by 4-hydroxyphenyl retinamide-induced NF-kappaB inhibition in acute myeloid leukemia cells. Cancer Chemother Pharmacol. 2020;86(2):257–66.

23. Morad SA, Davis TS, Kester M, Loughran TP, Jr., Cabot MC. Dynamics of ceramide generation and metabolism in response to fenretinide--Diversity within and among leukemia. Leuk Res. 2015;39(10):1071–8.

24. Potenza RL, Lodeserto P, Orienti I. Fenretinide in Cancer and Neurological Disease: A Two-Face Janus Molecule. Int J Mol Sci. 2022;23(13).

25. Takahashi N. Inhibitory Effects of Vitamin A and Its Derivatives on Cancer Cell Growth Not Mediated by Retinoic Acid Receptors. Biol Pharm Bull. 2022;45(9):1213–24.

26. Puduvalli VK, Yung WK, Hess KR, Kuhn JG, Groves MD, Levin VA, et al. Phase II study of fenretinide (NSC 374551) in adults with recurrent malignant gliomas: A North American Brain Tumor Consortium study. J Clin Oncol. 2004;22(21):4282–9.

27. Schneider BJ, Worden FP, Gadgeel SM, Parchment RE, Hodges CM, Zwiebel J, et al. Phase II trial of fenretinide (NSC 374551) in patients with recurrent small cell lung cancer. Investigational new drugs. 2009;27(6):571–8.

28. Garaventa A, Luksch R, Lo Piccolo MS, Cavadini E, Montaldo PG, Pizzitola MR, et al. Phase I trial and pharmacokinetics of fenretinide in children with neuroblastoma. Clinical cancer research : an official journal of the American Association for Cancer Research. 2003;9(6):2032–9.

29. Villablanca JG, Krailo MD, Ames MM, Reid JM, Reaman GH, Reynolds CP. Phase I trial of oral fenretinide in children with high-risk solid tumors: a report from the Children’s Oncology Group (CCG 09709). J Clin Oncol. 2006;24(21):3423–30.

30. Camerini T, Mariani L, De Palo G, Marubini E, Di Mauro MG, Decensi A, et al. Safety of the synthetic retinoid fenretinide: long-term results from a controlled clinical trial for the prevention of contralateral breast cancer. J Clin Oncol. 2001;19(6):1664–70.

31. Marcucci G, Silverman L, Eller M, Lintz L, Beach CL. Bioavailability of azacitidine subcutaneous versus intravenous in patients with the myelodysplastic syndromes. J Clin Pharmacol. 2005;45(5):597–602.

32. Kummar S, Gutierrez ME, Maurer BJ, Reynolds CP, Kang M, Singh H, et al. Phase I trial of fenretinide lym-x-sorb oral powder in adults with solid tumors and lymphomas. Anticancer research. 2011;31(3):961–6.

33. Mohrbacher AM, Yang AS, Groshen S, Kummar S, Gutierrez ME, Kang MH, et al. Phase I Study of Fenretinide Delivered Intravenously in Patients with Relapsed or Refractory Hematologic Malignancies: A California Cancer Consortium Trial. Clinical cancer research : an official journal of the American Association for Cancer Research. 2017;23(16):4550–5.

34. Badawi M, Chen X, Marroum P, Suleiman AA, Mensing S, Koenigsdorfer A, et al. Bioavailability Evaluation of Venetoclax Lower-Strength Tablets and Oral Powder Formulations to Establish Interchangeability with the 100 mg Tablet. Clinical Drug Investigation. 2022;42(8):657–68.

35. Zhao W, Sachsenmeier K, Zhang L, Sult E, Hollingsworth RE, Yang H. A New Bliss Independence Model to Analyze Drug Combination Data. J Biomol Screen. 2014;19(5):817–21.

36. Elcheva I, Brok-Volchanskaya V, Kumar A, Liu P, Lee JH, Tong L, et al. Direct induction of haematoendothelial programs in human pluripotent stem cells by transcriptional regulators. Nat Commun. 2014;5:4372.

37. Moorthi S, Burns TA, Yu GQ, Luberto C. Bcr-Abl regulation of sphingomyelin synthase 1 reveals a novel oncogenic-driven mechanism of protein up-regulation. FASEB J. 2018;32(8):4270–83.

38. Vu HL, Troubetzkoy S, Nguyen HH, Russell MW, Mestecky J. A method for quantification of absolute amounts of nucleic acids by (RT)-PCR and a new mathematical model for data analysis. Nucleic Acids Res. 2000;28(7):E18.

39. Tada H, Shiho O, Kuroshima K, Koyama M, Tsukamoto K. An improved colorimetric assay for interleukin 2. J Immunol Methods. 1986;93(2):157–65.

40. Lerdrup M, Hansen K. User-Friendly and Interactive Analysis of ChIP-Seq Data Using EaSeq. Methods Mol Biol. 2020;2117:35–63.

41. Vasu S, Kohlschmidt J, Mrózek K, Eisfeld AK, Nicolet D, Sterling LJ, et al. Ten-year outcome of patients with acute myeloid leukemia not treated with allogeneic transplantation in first complete remission. Blood Adv. 2018;2(13):1645–50.

42. Fujiwara Y, Browne CP, Cunniff K, Goff SC, Orkin SH. Arrested development of embryonic red cell precursors in mouse embryos lacking transcription factor GATA-1. Proc Natl Acad Sci U S A. 1996;93(22):12355–8.

43. Rodriguez P, Bonte E, Krijgsveld J, Kolodziej KE, Guyot B, Heck AJ, et al. GATA-1 forms distinct activating and repressive complexes in erythroid cells. EMBO J. 2005;24(13):2354–66.

44. Pevny L, Simon MC, Robertson E, Klein WH, Tsai SF, D’Agati V, et al. Erythroid differentiation in chimaeric mice blocked by a targeted mutation in the gene for transcription factor GATA-1. Nature. 1991;349(6306):257-60.

45. Welch JJ, Watts JA, Vakoc CR, Yao Y, Wang H, Hardison RC, et al. Global regulation of erythroid gene expression by transcription factor GATA-1. Blood. 2004;104(10):3136–47.

46. Phillips JD. Heme biosynthesis and the porphyrias. Molecular genetics and metabolism. 2019;128(3):164–77.

47. Zhang J, Wu K, Xiao X, Liao J, Hu Q, Chen H, et al. Autophagy as a regulatory component of erythropoiesis. Int J Mol Sci. 2015;16(2):4083–94.

48. Rahmaniyan M, Curley RW, Jr., Obeid LM, Hannun YA, Kraveka JM. Identification of dihydroceramide desaturase as a direct in vitro target for fenretinide. J Biol Chem. 2011;286(28):24754–64.

49. Decensi A, Formelli F, Torrisi R, Costa A. Breast cancer chemoprevention: Studies with 4-HPR alone and in combination with tamoxifen using circulating growth factors as potential surrogate endpoints. Journal of Cellular Biochemistry. 1993;53(S17G):226–33.

50. Bonanni B, Lazzeroni M, Veronesi U. Synthetic retinoid fenretinide in breast cancer chemoprevention. Expert Rev Anticancer Ther. 2007;7(4):423–32.

51. Agarwal SK, DiNardo CD, Potluri J, Dunbar M, Kantarjian HM, Humerickhouse RA, et al. Management of Venetoclax-Posaconazole Interaction in Acute Myeloid Leukemia Patients: Evaluation of Dose Adjustments. Clin Ther. 2017;39(2):359–67.

52. Leonard M, Brice M, Engel JD, Papayannopoulou T. Dynamics of GATA transcription factor expression during erythroid differentiation. Blood. 1993;82(4):1071–9.

53. Orienti I, Francescangeli F, De Angelis ML, Fecchi K, Bongiorno-Borbone L, Signore M, et al. A new bioavailable fenretinide formulation with antiproliferative, antimetabolic, and cytotoxic effects on solid tumors. Cell Death Dis. 2019;10(7):529.

54. Nguyen TH, Koneru B, Wei SJ, Chen WH, Makena MR, Urias E, et al. Fenretinide via NOXA Induction, Enhanced Activity of the BCL-2 Inhibitor Venetoclax in High BCL-2-Expressing Neuroblastoma Preclinical Models. Mol Cancer Ther. 2019;18(12):2270–82.

55. Corazzari M, Lovat PE, Oliverio S, Di Sano F, Donnorso RP, Redfern CP, et al. Fenretinide: a p53-independent way to kill cancer cells. Biochem Biophys Res Commun. 2005;331(3):810–5.

56. Stengel A, Kern W, Haferlach T, Meggendorfer M, Fasan A, Haferlach C. The impact of TP53 mutations and TP53 deletions on survival varies between AML, ALL, MDS and CLL: an analysis of 3307 cases. Leukemia. 2017;31(3):705–11.

57. Papaemmanuil E, Gerstung M, Bullinger L, Gaidzik VI, Paschka P, Roberts ND, et al. Genomic Classification and Prognosis in Acute Myeloid Leukemia. N Engl J Med. 2016;374(23):2209–21.

58. Rose D, Haferlach T, Schnittger S, Perglerova K, Kern W, Haferlach C. Subtype-specific patterns of molecular mutations in acute myeloid leukemia. Leukemia. 2017;31(1):11–7.

59. Cancer Genome Atlas Research N, Ley TJ, Miller C, Ding L, Raphael BJ, Mungall AJ, et al. Genomic and epigenomic landscapes of adult de novo acute myeloid leukemia. N Engl J Med. 2013;368(22):2059–74.

60. Fang H, Wang SA, Khoury JD, El Hussein S, Kim DH, Tashakori M, et al. Pure erythroid leukemia is characterized by biallelic TP53 inactivation and abnormal p53 expression patterns in de novo and secondary cases. Haematologica. 2022;107(9):2232–7.

61. Iacobucci I, Qu C, Varotto E, Janke LJ, Yang X, Seth A, et al. Modeling and targeting of erythroleukemia by hematopoietic genome editing. Blood. 2021;137(12):1628–40.

62. Lal R, Lind K, Heitzer E, Ulz P, Aubell K, Kashofer K, et al. Somatic TP53 mutations characterize preleukemic stem cells in acute myeloid leukemia. Blood. 2017;129(18):2587–91.

63. Rucker FG, Schlenk RF, Bullinger L, Kayser S, Teleanu V, Kett H, et al. TP53 alterations in acute myeloid leukemia with complex karyotype correlate with specific copy number alterations, monosomal karyotype, and dismal outcome. Blood. 2012;119(9):2114–21.

64. Pan X, Ohneda O, Ohneda K, Lindeboom F, Iwata F, Shimizu R, et al. Graded levels of GATA-1 expression modulate survival, proliferation, and differentiation of erythroid progenitors. J Biol Chem. 2005;280(23):22385–94.

65. Yu YL, Chiang YJ, Chen YC, Papetti M, Juo CG, Skoultchi AI, et al. MAPK-mediated phosphorylation of GATA-1 promotes Bcl-XL expression and cell survival. J Biol Chem. 2005;280(33):29533–42.

66. Clifford JL, Menter DG, Wang M, Lotan R, Lippman SM. Retinoid receptor-dependent and -independent effects of N-(4-hydroxyphenyl)retinamide in F9 embryonal carcinoma cells. Cancer Res. 1999;59(1):14–8.

67. Ponzoni M, Bocca P, Chiesa V, Decensi A, Pistoia V, Raffaghello L, et al. Differential effects of N-(4-hydroxyphenyl)retinamide and retinoic acid on neuroblastoma cells: apoptosis versus differentiation. Cancer Res. 1995;55(4):853–61.

68. Oridate N, Suzuki S, Higuchi M, Mitchell MF, Hong WK, Lotan R. Involvement of reactive oxygen species in N-(4-hydroxyphenyl)retinamide-induced apoptosis in cervical carcinoma cells. J Natl Cancer Inst. 1997;89(16):1191–8.

69. Kitareewan S, Spinella MJ, Allopenna J, Reczek PR, Dmitrovsky E. 4HPR triggers apoptosis but not differentiation in retinoid sensitive and resistant human embryonal carcinoma cells through an RARgamma independent pathway. Oncogene. 1999;18(42):5747–55.

70. Ruvolo VR, Karanjeet KB, Schuster TF, Brown R, Deng Y, Hinchcliffe E, et al. Role for PKC delta in Fenretinide-Mediated Apoptosis in Lymphoid Leukemia Cells. J Signal Transduct. 2010;2010:584657.

71. Yang H, Bushue N, Bu P, Wan YJ. Induction and intracellular localization of Nur77 dictate fenretinide-induced apoptosis of human liver cancer cells. Biochem Pharmacol. 2010;79(7):948–54.

72. Yang H, Zhan Q, Wan YJ. Enrichment of Nur77 mediated by retinoic acid receptor beta leads to apoptosis of human hepatocellular carcinoma cells induced by fenretinide and histone deacetylase inhibitors. Hepatology. 2011;53(3):865–74.

73. Tiwari M, Kumar A, Sinha RA, Shrivastava A, Balapure AK, Sharma R, et al. Mechanism of 4-HPR-induced apoptosis in glioma cells: evidences suggesting role of mitochondrial-mediated pathway and endoplasmic reticulum stress. Carcinogenesis. 2006;27(10):2047–58.

74. De Maria R, Zeuner A, Eramo A, Domenichelli C, Bonci D, Grignani F, et al. Negative regulation of erythropoiesis by caspase-mediated cleavage of GATA-1. Nature. 1999;401(6752):489–93.

