## Supplementary material all for "Fenretinide targets GATA1 to induce cytotoxicity in GATA1 positive Acute Erythroid and Acute Megakaryoblastic Leukemic cells"

**Supplementary Materials**

**Figure Legends**

**Figure S1: *GATA1* expression in Chronic Myeloid Leukemia (CML) cell lines**. *GATA1* expression across CML cell lines reveal high GATA1 expression, according to data from Cancer Cell Line Encyclopedia (CCLE).

**Figure S2: 4-HPR treatment of GATA1 negative HL-60 cells and ATRA treatment of OCIM1 cells have no impact on cell number.** (**A**) HL-60 cells, classified as M2/M3 AML, were treated with 0, 0.5, 1, 1.5, and 2µM of 4-HPR which had no impact on cell number over 48 hours of treatment. Reflective of three independent experiments. **(B)** OCIM1 cells were treated with 0-10µM of all-trans retinoic acid (ATRA). ATRA does not significantly impact cell number up to 48 hours after treatment. Reflective of two independent experiments.

**Figure S3: 4-HPR treatment of OCIM1 cells reduces *GATA1* mRNA levels.** OCIM1 cells were treated with 0, 200, 350, 500, and 750nM of 4-HPR. mRNA expression of *GATA1* was measured by qRT-PCR and normalized to *18S.* **p<0.005

**Figure S4: High doses of 4-HPR treatment in combination with Azacytidine and Venetoclax.** (**A**) The ability of 4-HPR to decrease viability of OCIM1 cells is represented by percent inhibition measured by MTT assay. Concentrations of Azacytidine (Aza) pre-treatment at 0, 2, 5, 7µM (with and without additional 0.5µM of Aza maintained in media) and Venetoclax (Veneto) at 0, 1, 2, and 3µM as indicated, were tested in the presence of 0.5, 1, and 2µM of 4-HPR for sensitization to Aza+Veneto. (**B**) Synergy scores (δ – score) of 4-HPR (0.5, 1, and 2µM) with Aza and Veneto, with or without maintenance of 0.5µM Aza in the media after 48 hours. δ – scores were determined by SynergyFinder (synergyfinder.fimm.fi), according to Bliss independence, where δ – score > 10 indicates synergism, and δ – score > -10 indicates antagonism.

### Tables

**Supplementary Table 1:** *GATA1* over-expression construct primers

| Primer | Sequence (5' to 3') | |
|---|---|---|
| *GATA1* | Forward (with Kpn-I site) | CAATGGTACCATG<br>GAGTTCCCTG |
| | Reverse (with Xba-I site) | CACTCTAGATCATG<br>AGCTGA |

**Supplementary Table 2:** *GATA1* cDNA sequence for over-expression.

ATGGAGTTCCCTGGCCTGGGGTCCCTGGGGACCTCAGAGCCCCTCCCCCAGTTTGTGGATCCTGCTCTGGTGTCCTCCACACCAGAATCAGGGGTTT
TCTTCCCCTCTGGGCCTGAGGGCTTGGATGCAGCAGCTTCCTCCACTGCCCCGAGCACAGCCACCGCTGCAGCTGCGGCACTGGCCTACTACAGGGA
CGCTGAGGCCTACAGACACTCCCCAGTCTTTCAGGTGTACCCATTGCTCAACTGTATGGAGGGGATCCCAGGGGGCTCACCATATGCCGGCTGGGCC
TACGGCAAGACGGGGCTCTACCCTGCCTCAACTGTGTGTCCCACCCGCGAGGACTCTCCTCCCCAGGCCGTGGAAGATCTGGATGGAAAAGGCAGCA
CCAGCTTCCTGGAGACTTTGAAGACAGAGCGGCTGAGCCCAGACCTCCTGACCCTGGGACCTGCACTGCCTTCATCACTCCCTGTCCCCAATAGTGC
TTATGGGGGCCCTGACTTTTCCAGTACCTTCTTTTCTCCCACCGGGAGCCCCCTCAATTCAGCAGCCTATTCCTCTCCCAAGCTTCGTGGAACTCTC
CCCCTGCCTCCCTGTGAGGCCAGGGAGTGTGTGAACTGCGGAGCAACAGCCACTCCACTGTGGCGGAGGGACAGGACAGGCCACTACCTATGCAACG
CCTGCGGCCTCTATCACAAGATGAATGGGCAGAACAGGCCCCTCATCCGGCCCAAGAAGCGCCTGATTGTCAGTAAACGGGCAGGTACTCAGTGCAC
CAACTGCCAGACGACCACCACGACACTGTGGCGGAGAAATGCCAGTGGGGATCCCGTGTGCAATGCCTGCGGCCTCTACTACAAGCTACACCAGGTG
AACCGGCCACTGACCATGCGGAAGGATGGTATTCAGACTCGAAACCGCAAGGCATCTGGAAAAGGGAAAAAGAAACGGGGCTCCAGTCTGGGAGGCA
CAGGAGCAGCCGAAGGACCAGCTGGTGGCTTTATGGTGGTGGCTGGGGGCAGCGGTAGCGGGAATTGTGGGGAGGTGGCTTCAGGCCTGACACTGGG
CCCCCCAGGTACTGCCCATCTCTACCAAGGCCTGGGCCCTGTGGTGCTGTCAGGGCCTGTTAGCCACCTCATGCCTTTCCCTGGACCCCTACTGGGC
TCACCCACGGGCTCCTTCCCCACAGGCCCCATGCCCCCCACCACCAGCACTACTGTGGTGGCTCCGCTCAGCTCATGA

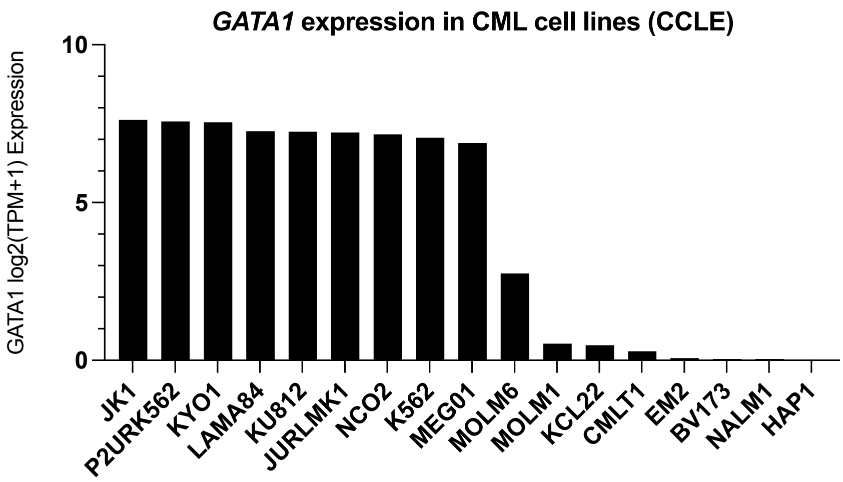


Figure S1: GATA1 expression in Chronic Myeloid Leukemia cell lines.

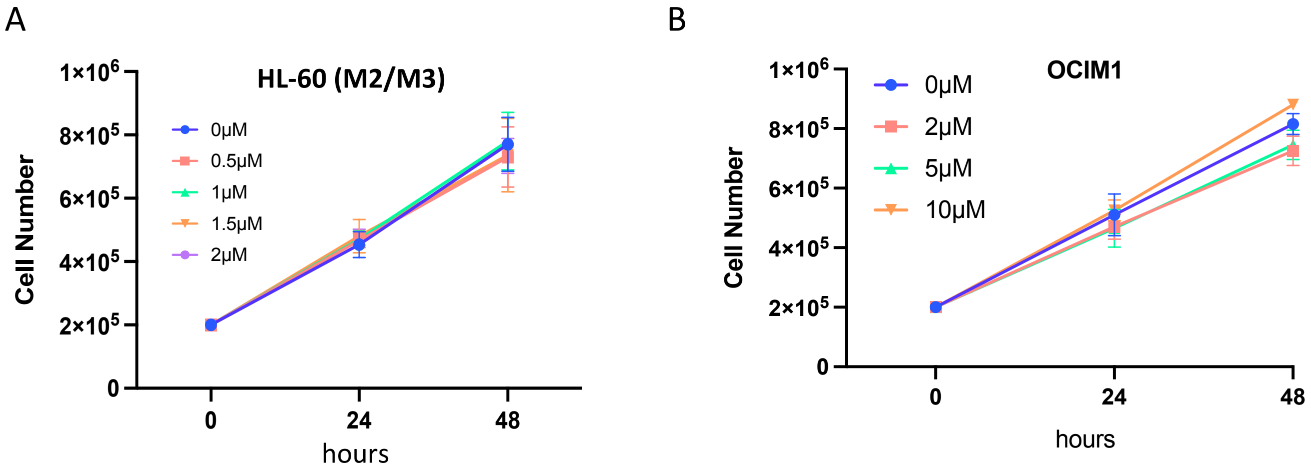


Figure S2: 4-HPR treatment of GATA1 negative HL-60 cells and ATRA treatment of OCIM1 cells have no impact on cell number.

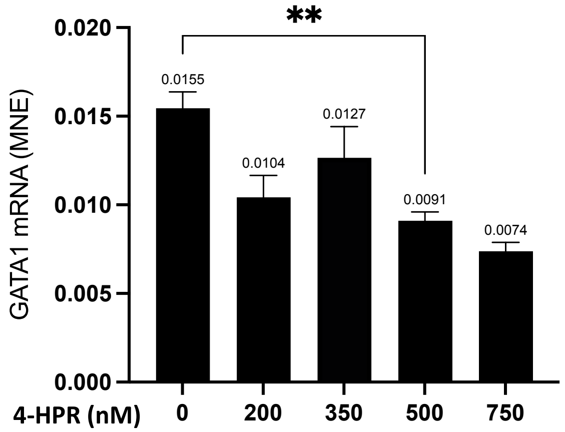


Figure S3: 4-HPR treatment of OCIM1 cells reduces *GATA1* mRNA levels

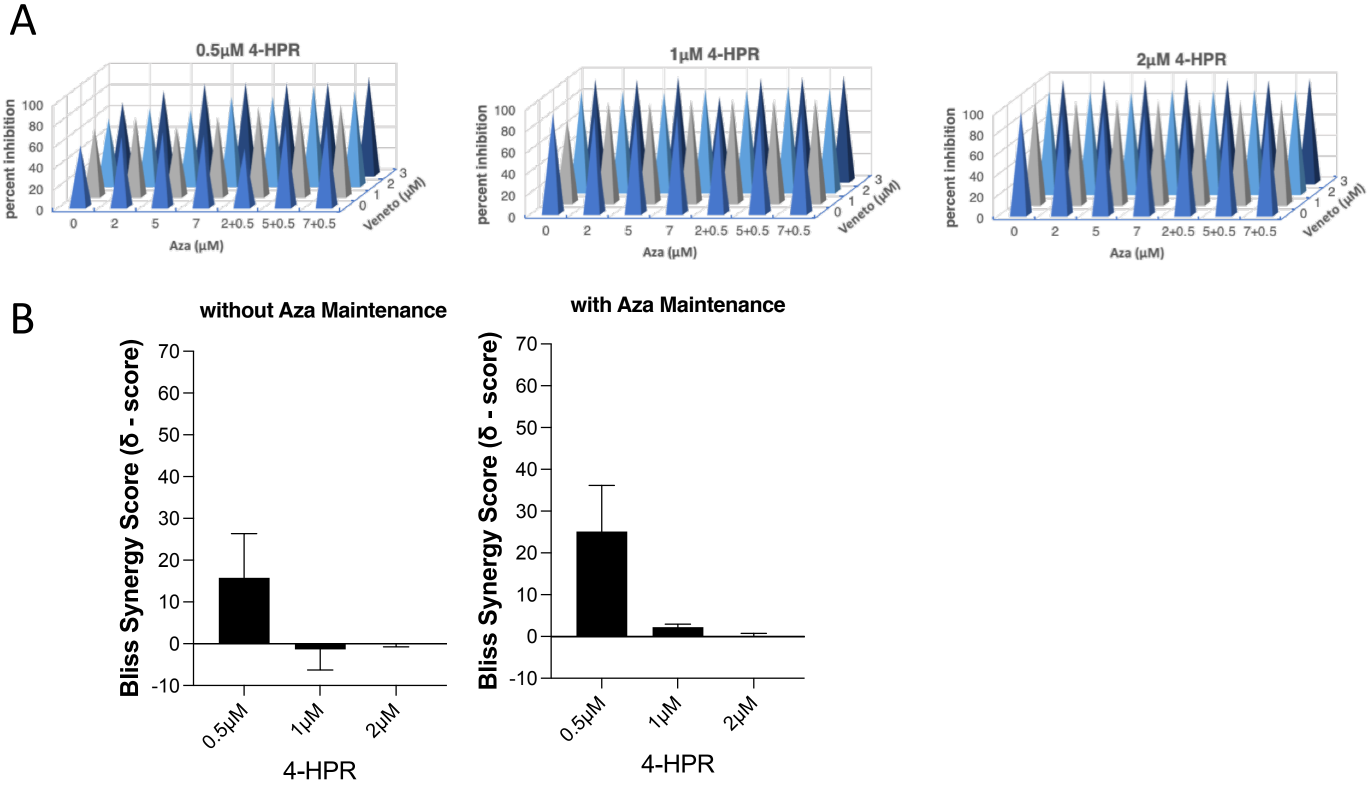


Figure S4: High doses of 4-HPR treatment in combination with Azacytidine and Venetoclax
